# Population-scale chemical response revealed by a barcoded yeast collection

**DOI:** 10.64898/2025.12.03.691428

**Authors:** Abhishek Dutta, Marion Garin, Victor Loegler, Gauthier Brach, Anne Friedrich, Mami Yoshimura, Hiroyuki Hirano, Hiroyuki Osada, Charles Boone, Yoko Yashiroda, Jing Hou, Joseph Schacherer

## Abstract

Natural genetic variation shapes how microbial populations adapt to environmental and chemical challenges, but scalable approaches to map genotype-phenotype relationships across diverse genetic backgrounds remain limited. Here, we developed a systematically barcoded collection of 520 *Saccharomyces cerevisiae* natural isolates that captures the ecological, geographical and genetic diversity of the species. Using pooled barcode sequencing, we profiled fitness responses to over 600 bioactive and natural compounds, revealing broader and more polarized bioactivity than the standard yeast gene-deletion collection. Fitness-based clustering defined six major compound groups with reproducible, population-structured sensitivity patterns. Genome-wide association analysis identified significant genetic variants across 107 compounds, linking natural polymorphisms to chemical responses and involving genes in genome maintenance, ribosome biogenesis, vesicular trafficking and stress tolerance. Together, our barcoded natural population provides a scalable framework for chemical-genetic screening, enabling systematic dissection of how genetic diversity shapes microbial fitness and adaptation.

## Introduction

Phenotypic traits within a species are highly variable. Populations harbour natural genetic diversity and continually acquire mutations that shape the phenotypic landscape ^1–4^. This diversity fuels evolution and determines how populations adapt to new environments, yet the genetic basis of phenotypic variation remains only partly understood ^5–9^.

Microbial systems vividly illustrate this complexity. In bacteria, genetic diversity drives stress tolerance, antibiotic resistance, and biofilm formation ^10–12^. In viruses, intra-host diversity enables immune evasion and drug resistance ^13,14^. In yeasts such as *Candida albicans* and *Saccharomyces cerevisiae*, variation in morphology, metabolism, and stress response underlies virulence, ecological specialization, and diversification ^15–20^. Together, such genetic and phenotypic variation shapes fitness and evolutionary trajectories ^21–23^. Similar dynamics drive tumour evolution, where genetic diversity fosters phenotypic heterogeneity, adaptation, and therapy resistance ^24–26^.

Mapping genotype-phenotype relationships is fundamental to understanding the genetic architecture of complex traits and identifying causal variants underlying disease and drug resistance ^27–29^. Despite extensive genomic resources, accurately linking genetic variation to phenotypic variation remains difficult. Comprehensive mapping requires large, reproducibly phenotyped populations, which is still a major bottleneck in many systems ^17,30,31^. High-throughput functional genomics, particularly barcode sequencing (Bar-seq), has transformed phenotypic profiling at scale. Unlike traditional strain-by-strain assays limited by throughput and reproducibility, Bar-seq enables pooled fitness measurements for thousands of uniquely tagged genotypes in a single experiment ^32–36^. This innovation revolutionized chemical-genetic screening in *S. cerevisiae* deletion mutant collections, facilitating large-scale functional annotation and the discovery of novel bioactive compounds ^35,37–41^. However, these approaches remain largely restricted to loss-of-function variants in a few reference strains and fail to capture the allelic, structural, and regulatory diversity found in natural populations ^17,42,43^. Consequently, much of the biologically relevant genotype-phenotype landscape remains unexplored.

Wild and domesticated populations of *S. cerevisiae* display remarkable ecological, geographical, and genetic diversity, encompassing hundreds of sequenced isolates from both natural and human-associated environments ^17,44–47^. This diversity makes *S. cerevisiae* an outstanding model for studying how genetic variation shapes phenotypic diversity. Coupled with extensive genomic resources, genome-wide association studies (GWAS) have enabled the identification of naturally segregating variants that influence cellular responses across diverse environments^17,30,48–51^. Exploiting this natural variation helps to reveal the regulatory and functional diversity that underlies population-level adaptation to environmental and chemical challenges.

Here, we developed a systematically barcoded collection of 520 natural *S. cerevisiae* isolates representing the global ecological and genetic diversity of the species. Benchmarking confirmed that this collection recapitulates the phenotypic distributions and associations observed in the original population, validating its suitability for pooled fitness profiling. Screening approximately 600 bioactive and natural compounds showed that the barcoded population exhibits broader and stronger bioactivity than a drug-sensitized laboratory deletion collection, underscoring its enhanced responsiveness to chemical perturbations ^40^. Fitness-based clustering of compound responses identified six major groups with shared modes of action and reproducible, population-structured sensitivity patterns. Genome-wide association analyses (GWAS) revealed 46 genes linked to responses across 107 compounds, connecting natural polymorphisms to chemical phenotypes. By integrating natural genetic diversity with large-scale chemical profiling, this work establishes a powerful and scalable framework for systematic phenotyping across hundreds to thousands of compounds. The barcoded collection enables detailed analysis of adaptive traits and compound responses, complementing traditional deletion-based approaches by capturing naturally evolved variation in its native genomic context. Together, these advances expand the phenotypic landscape of *S. cerevisiae* and pave the way toward a high-resolution genotype-phenotype map in yeast.

## Results

### Generation of a barcoded collection of natural *S. cerevisiae* isolates

To expand the phenotypic landscape and enable pooled chemical-genetic screening across natural diversity, we set out to barcode the 1,011 natural *S. cerevisiae* isolates ^17^. Using a construct designed for integration at the *HO* locus, each barcode consisted of a unique 20-nucleotide sequence linked to a selectable marker, allowing individual strains to be tracked within mixed populations (Figure 1A).

**Figure 1.**
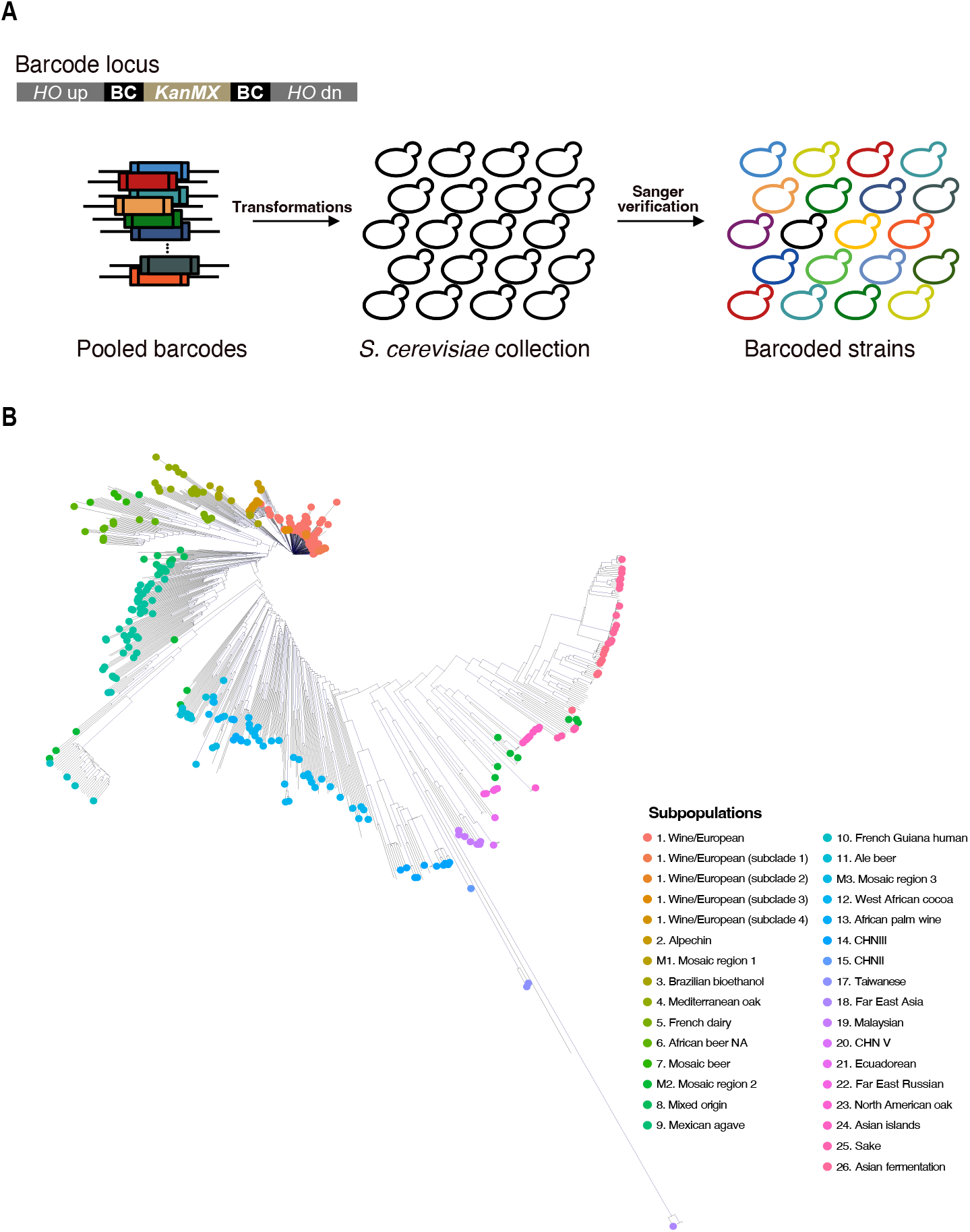
Barcoding strategy and genetic diversity of the yeast strain collection. **A**. Schematic representation of the barcode locus and the barcoding of the *S. cerevisiae* strain collection. **B**. Neighbour-joining tree built using the biallelic SNPs representing diversity of the barcoded strains in context of the 1,011 *S. cerevisiae* population.

Following multiple rounds of transformation, barcode integration was confirmed in more than half of the isolates. Integration failed in a subset of strains, likely due to polymorphisms in the *HO* locus or natural variability in transformation efficiency among isolates. In total, we successfully barcoded 538 isolates spanning 53.2% of the collection (Table S1). Sanger sequencing confirmed that the majority of these isolates (n=524) carries a 20-nucleotide barcodes as designed, whereas a small fraction (n=14) carries slightly shorter or longer barcodes (14–28 nt), likely reflecting errors during barcode synthesis or PCR-based extension. Despite the slight size variation, the high average levenshtein edit distance of 12 (range 3–18) confirmed that the barcodes were sufficiently diverse to uniquely identify each strain (Figure S1A). The mean GC content was 49% (range 15–85%), with 95% of barcodes falling within the optimal amplification range of 30–70% GC (Figure S1B). Conservative quality thresholds (edit distance >4 and GC content between 30% and 70%) yielded a high-confidence barcoded collection of 520 isolates, representing 51.4% of the original population (Figure 1B).

To determine whether the barcode integration affected strain fitness, we compared the growth phenotypes of barcoded isolates with their non-barcoded counterparts across two environments (rich YPD and minimal SC). Fitness values were strongly correlated between barcoded and non-barcoded strains (Spearman’s ρ = 0.81-0.83; p < 2 × 10^-16^), with a median fitness ratio of 1.008 and no significant deviation from equivalence (Wilcoxon p = 0.71-0.83), confirming that the barcode integration did not impact strain fitness across tested environments (Figures S2A-B). These 520 barcoded isolates formed a representative subset of the original 1,011 population, mirroring its genetic, ecological, and geographic diversity (π = 3.6 × 10^−3^), making it a powerful resource for population-scale chemical-genetic and genome-wide association studies (Figures 2B–D) ^17,49^.

**Figure 2.**
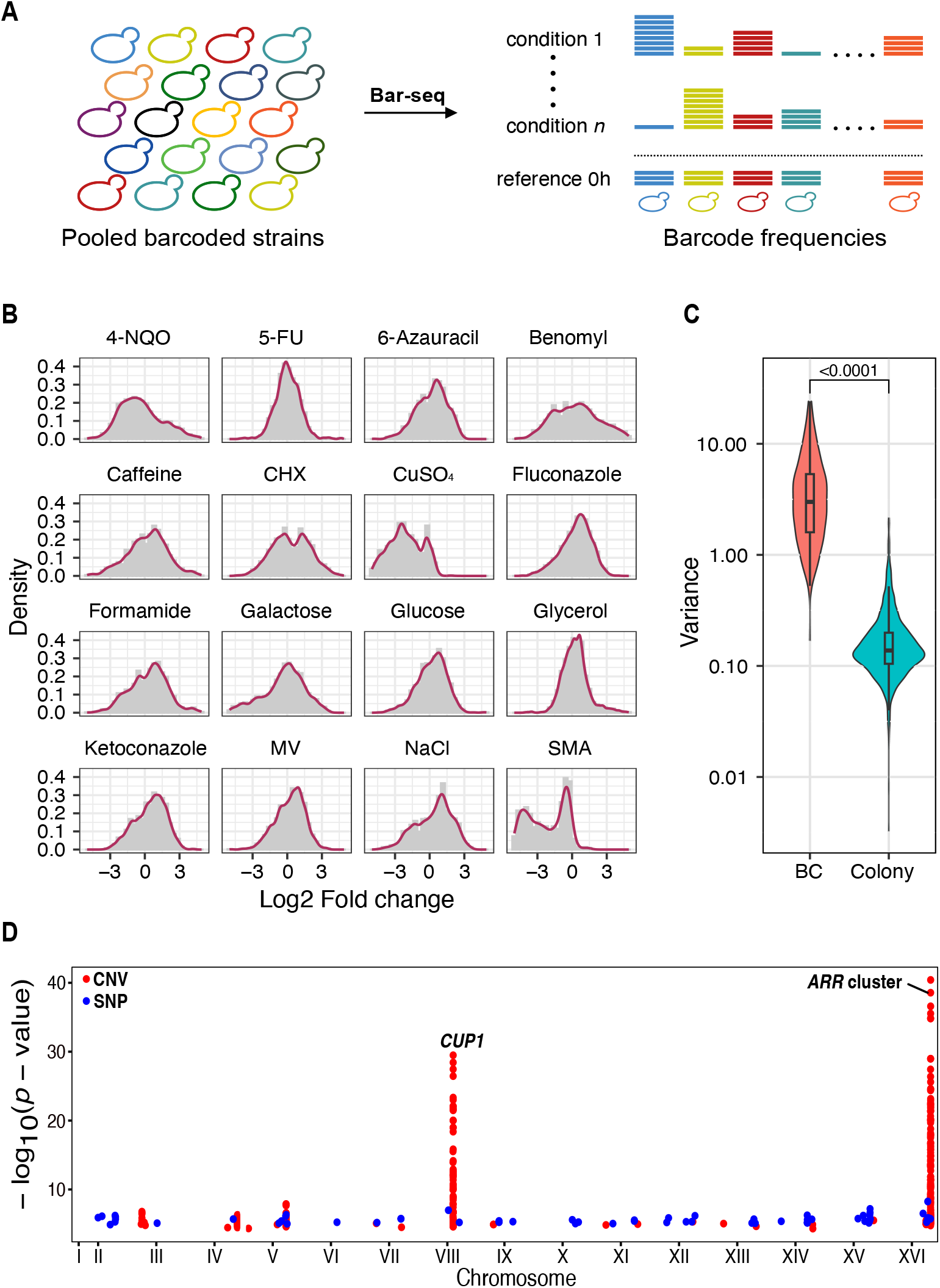
Benchmarking the barcoded collection. **A**. Schematic representation of the barcode-sequencing (Bar-seq) experimental strategy. **B**. Representative fitness distributions (log2 fold change) of the barcoded isolates across the 16 compounds tested. **C**. Violin-plot representing the differences in the phenotypic variance of the 495 barcoded isolates via the barcode sequencing strategy and colony-size based phenotyping. **D**. Manhattan-plot representing only the 111 significant variants detected in Bar-seq GWAS. Solid red speres represent CNVs and solid blue circles represent SNPs across the 16 chromosomes.

### Benchmarking the barcoded collection

To evaluate the performance and utility of the barcoded collection, we conducted a proof-of-concept experiment using the Bar-seq strategy across 45 conditions encompassing 16 compounds targeting diverse cellular processes, including transcription, translation, carbon metabolism, and DNA repair (Figure 2A; Table S2). These conditions were selected because the original 1,011-collection and its derivatives had been extensively phenotyped by colony-size on solid media, providing an ideal benchmark to compare our Bar-seq results with colony-based phenotypes using the same subset of strains ^17,52^.

A master pool of 495 barcoded isolates was created after excluding strains with genome-encoded killer viruses or flocculation phenotypes to prevent confounding interactions (Table S1) ^53^. Pooled competition assays were performed in 200 µL volumes supplemented with test compounds, and cells were harvested at 0h and 24h timepoints for DNA extraction, library preparation, and barcode sequencing. Barcode sequencing detected 490 of the 495 unique strains at the 0h timepoint and 430– 444 strains across conditions at 24h after applying conservative thresholds (see Materials and Methods). At 0h, barcode fractions were evenly distributed, with an average relative abundance of 0.22% (range 0.01–1.8%), while at 24 h the distributions broadened substantially, reflecting differential strain growth profiles under each condition (Figure 2B; Table S2). Fitness values for each strain were estimated as the log_2_ fold change in relative barcode abundance from 0 h to 24 h, providing a measure of relative growth within the pooled population. Fitness estimates were highly reproducible among replicates (Pearson’s r = 0.82–0.99) (Table S2). The fitness distributions across conditions closely mirrored those from the full 1,011 collection, confirming that the barcoded subset preserved the phenotypic diversity of the entire population (Figure 2B). We compared the diversity of phenotypes obtained from colony size-based assays with those derived from barcode-based measurements and found that phenotypic variance was, on average, an order of magnitude higher in the Bar-seq data (Figure 2C). This highlights the ability of bar-seq approach to capture subtle quantitative effects.

To connect these phenotypes to their genetic basis, we performed genome-wide association studies (GWAS) using a high-density genotype matrix of 91,317 SNPs and 700 CNVs ^17,49^. Our GWAS identified 111 genetic variants (93 SNPs and 18 CNVs) significantly associated with fitness under 13 different compounds (Figure 2D; Table S3). CNVs explained a greater proportion of variance than SNPs (median 30.2% vs. 22.4%) (Figure S3). The amplification of the *CUP1* and *ARR* gene cluster were strongly associated with resistance to CuSO_4_ (p = 3.9 × 10^−30^) and sodium meta-arsenite (p = 3.6 × 10^−22^), respectively, together accounting for over 45% of the observed phenotypic variance. These GWAS results were highly comparable to previous results obtained using colony-based growth phenotypes, in particular for phenotypes driven by common, high effect variants.

Together, these results demonstrate that the barcoded collection effectively captures the phenotypic diversity and genetic associations of the original population. Importantly, the small assay volumes and scalability of the Bar-seq strategy establish it as a powerful high-throughput platform for phenotyping across hundreds or thousands of conditions, thereby expanding phenotypic landscape at the species level.

### Natural variation broadens the fitness-response landscape

Leveraging our robust barcoded collection, we deployed this high-throughput platform to screen a library of 633 naturally derived compounds from the RIKEN Natural Product Depository (NPDepo). This library consists primarily of purified natural products or their derivatives. A full description of the compounds and their structures can be found in Table S4.

We performed pooled competition assays, barcode sequencing, and phenotypic quantification using the methods established in our proof-of-concept experiments. We detected 475 (~96%) strains at the 0h timepoint (Table S5). At the 24h timepoint, an average of 400 strains were detected across most conditions (Figure S4; Table S6). Following initial screening, 178 compounds were excluded. These included 120 compounds (~19%) that showed complete growth inhibition after 24 hours and 58 compounds that did not pass our data quality control criteria (see Materials and methods) (Figure 3A). Strain fitness was defined as the log2 fold-change in barcode abundance between the 24h and 0h timepoints.

**Figure 3.**
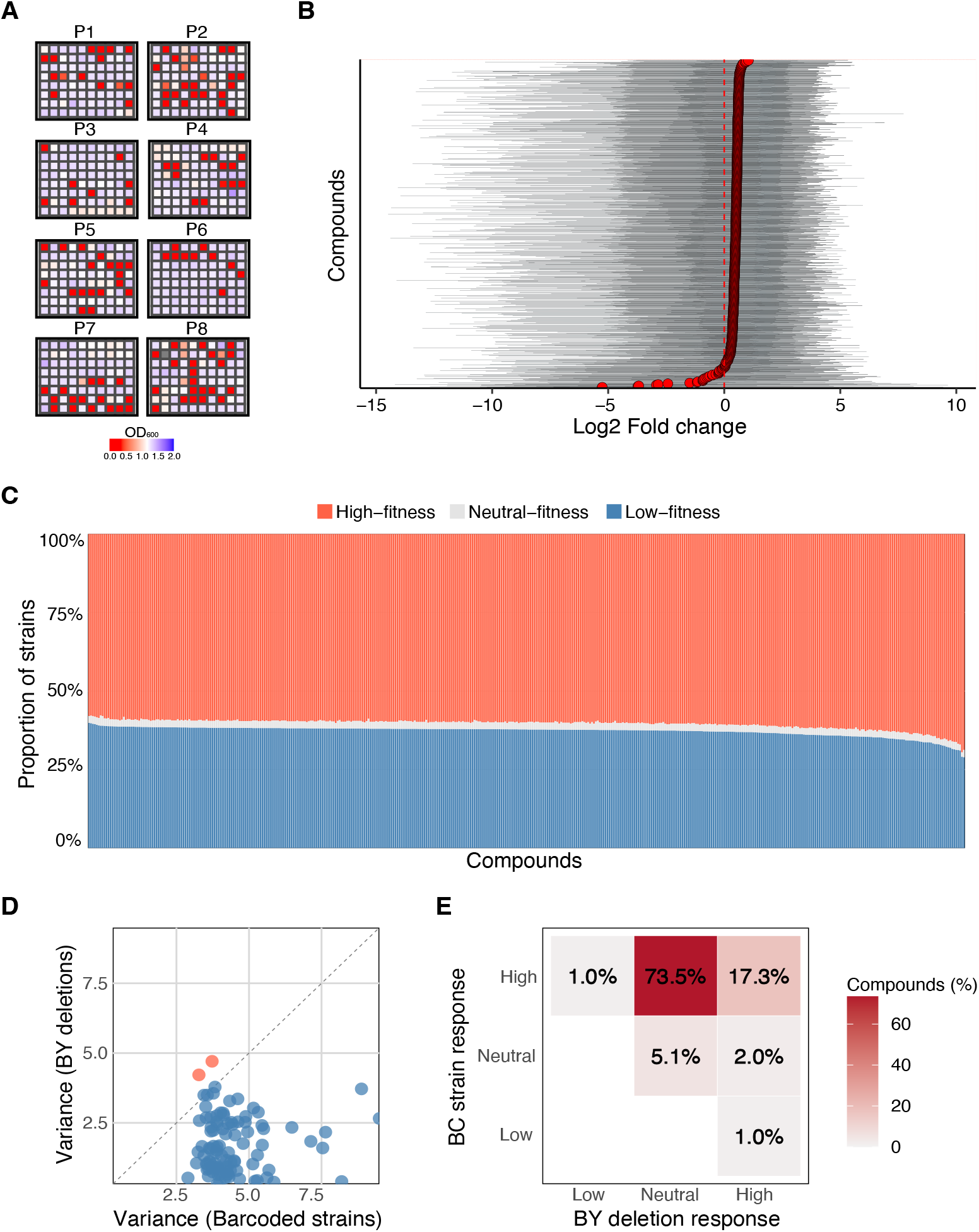
Large-scale compound-response profiling of barcoded yeast population. **A**. Blank-corrected OD_600_ values of the barcode pools across compound library after 24 hours. The red wells denote the compounds that completely inhibited growth. **B**. Violin-plot representing the fitness values of the strains in the barcoded pool across 455 compounds. The solid red dot indicates mean pool fitness in the compound and the dashed red line indicates Log2-fold change=0. **C**. Marplot representing the fitness-response profile proportions of the individual strains in the pool across 455 compounds based on our binomial classification. Red, grey and blue indicate the proportions of high-fitness, neutral-fitness and low-fitness strains respectively. **D**. Scatter-plot showing the correlation between fitness variance of barcoded and BY deletion strains across 98 shared compounds (Spearman’s ρ = –0.05). Blue circles denote compounds with higher variance in the barcoded strains (96/98), and red circles denote compounds with higher variance in the BY deletion strains (2/98). **E**. Heatmap comparing the fitness response profiles of the Barcoded and BY deletion strains across the 98 shared compounds.

The fitness of strains in the barcoded pool across the remaining 455 compounds was highly variable. The overall fitness distribution was asymmetric, with a pronounced shift away from neutrality (Wilcoxon test, p < 2 × 10^−16^) (Figure 3B, S5; Table S6), indicating that a substantial subset of strains consistently exhibited non-random responses across multiple chemical perturbations. To classify individual strain responses, we applied a binomial framework that tested whether the proportion of positive versus negative fold-changes for each strain deviated from the null expectation (p = 0.5). Resulting p-values were FDR-corrected. We designated strains with significantly enriched positive responses (FDR < 0.05) as high-fitness (n=248), those with significantly enriched negative responses as low-fitness (n=170), and the remainder as neutral-fitness (n=26) (Table S7). This classification revealed that the majority of strains exhibited consistent directional responses across the compound library (Figure 3C). In contrast, the neutral-fitness group (~6% of strains) showed reduced variance and lower net fitness effects across compounds (Table S7). These results highlight that the barcoded strain population is broadly responsive to the compound library, revealing extensive phenotypic diversity in chemical tolerance.

Previously, *S. cerevisiae* deletion mutant collections have been extensively used to profile compound libraries and map chemical-genetic interaction landscapes ^37,40,54–56^. To place our findings in context, we compared the fitness responsiveness landscape of our barcoded natural strain population with that of the hyper sensitized reference BY gene deletion collection screened against a similar compound library ^40^. In total, 98 compounds were shared between Piotrowski et al ^40^. and this study. In addition, it is important to note that the deletion collection was screened at a uniform compound concentration of 10 µg mL^−1^, whereas our barcoded collection was screened at 100 µM, which correspond to an average of ~8 µg mL^−1^, with 80% of compounds below 10 µg mL^−1^ therefore at a lower concentration than that was used for the deletion collection previously (Figure S6).

Applying our binomial framework to the BY deletion mutant dataset revealed that 62% showed high-fitness, 30% showed neutral fitness and only 8% of strains were classified low-fitness. In contrast, only 6% of strains in our barcoded population were classified as neutral, indicating that natural genetic variation yields a more polarized and responsive fitness landscape. Across the 98 compounds shared between the two datasets, we quantified the phenotypic variance of each strain collection and found that the barcoded pool displayed markedly higher variance for 96 of the 98 compounds, while the deletion collection showed higher variance in only two (Figure 3D; Table S8). Comparing compound-wise classifications further revealed that 72 (~73.5%) of the shared compounds exhibited a significant high-fitness bias among barcoded strains but were neutral in the deletion dataset, whereas only 17 compounds (~17.3%) were high-fitness in both, and reciprocal or negative biases were rare (~1%) (Figure 3E).

Together, these results highlight that the natural diversity within the barcoded collection show significant bioactivity toward the compound library. Remarkably, compared to deletion mutants in the hyper-sensitized reference BY background, a broader range of chemical responsiveness in is evident in the natural population, providing a potentially better platform to profile the biological functions for large natural compound libraries.

### Fitness-based functional profiling of the natural compound library

We aimed to profile the natural compound library based on the fitness response patterns of the barcoded strains to delineate it into biologically meaningful clusters. Such an approach enables the discovery of shared physiological targets and reveals how natural genetic diversity shapes chemical sensitivity a crucial step for linking compound activity to underlying biological mechanisms ^40^. To uncover global structure of the compounds in the strain fitness response space, we computed pairwise correlations between all strain fitness profiles across compounds. The resulting similarity matrix partitioned these compounds into discrete groups with highly similar strain-level response signatures (Figure 4A). Hierarchical clustering of this matrix resolved into six well-supported compound clusters, each defined by coherent and reproducible strain fitness profiles (Figure 4B; Table S9). These clusters capture compounds that elicit coherent phenotypic effects across the strain panel, suggesting convergence on related physiological pathways or stress responses.

**Figure 4.**
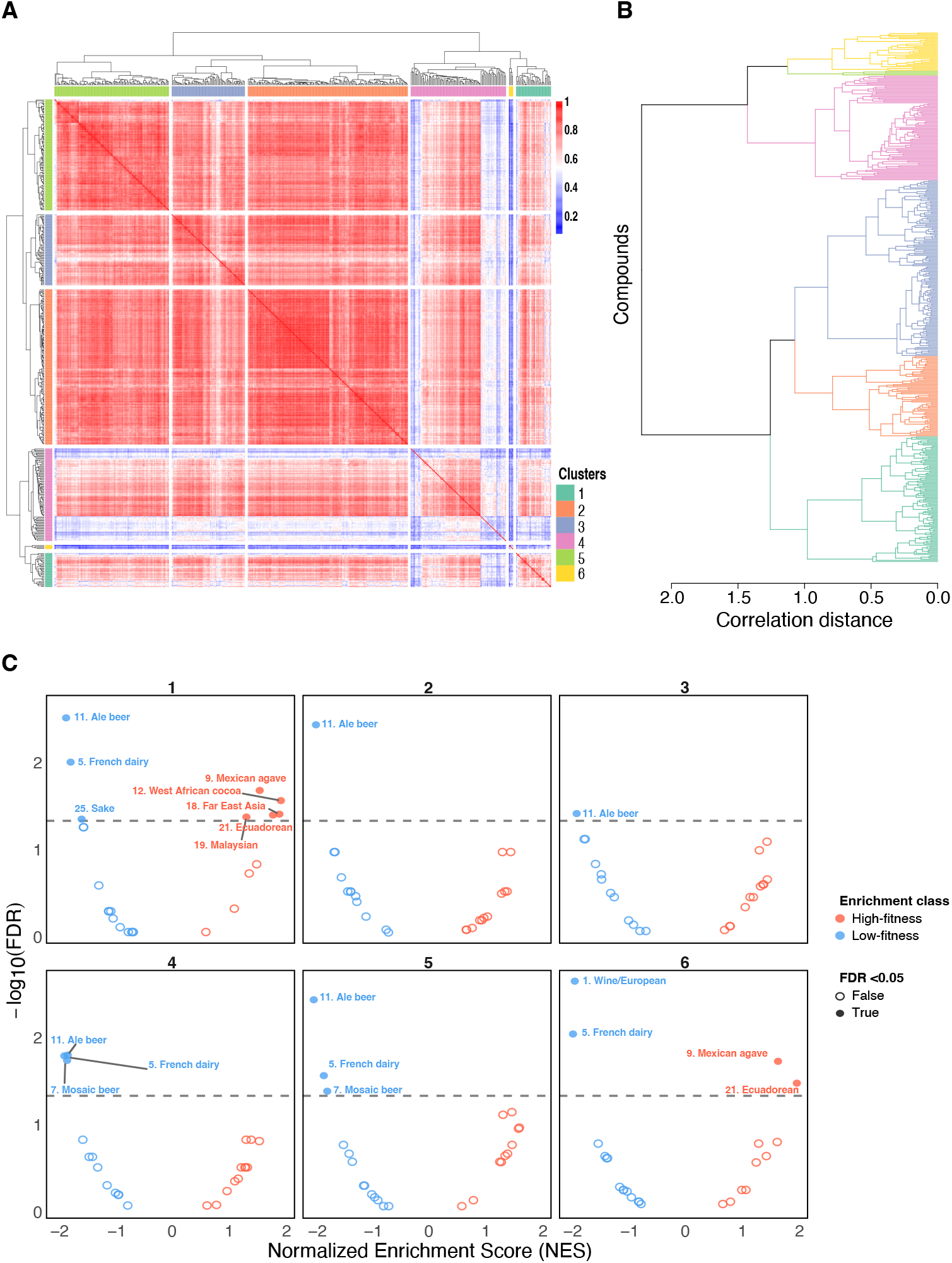
Clustering of compounds by strain-level fitness. **A**. Compound–compound correlation matrix based on strain fitness (log_2_ fold-change) profiles. Blocks of high correlation (red) indicate groups of compounds eliciting similar fitness responses across the strain panel, revealing strong modular organization within the dataset. **B**. Hierarchical clustering dendrogram of compounds derived from the correlation matrix, defining six robust clusters (colored branches) that represent distinct modes of compound-induced response similarity. **C**. Subpopulation enrichment analysis (GSEA-style) showing the distribution of high- and low-fitness classes across the six compound clusters. Each cluster exhibits a characteristic pattern of subpopulation enrichment, demonstrating that the compound clusters are differentiated by reproducible, subpopulation-specific fitness response signatures.

We tested whether these compound-defined groups corresponded to underlying population structure by performing a GSEA-like subpopulation-based enrichment analysis ^57^. Our analysis revealed that each compound cluster displayed a distinct pattern of subpopulation enrichment among high- and low-fitness responders (Figure 4C). This indicates that the six clusters are structured not only by overall fitness similarity but also by consistent, lineage-specific response biases. For example, Ale beer and French dairy subpopulations were consistently enriched among low-fitness responders across several clusters, whereas Mexican agave, Ecuadorian, and West African cocoa subpopulations were enriched in the high-fitness class.

To identify the individual strains driving these patterns, we quantified per-cluster strain signatures, which represent the mean normalized fitness of each strain within a cluster (Table S10A). These signatures revealed cluster-specific marker strains that define each group’s unique response profile. In parallel, we used a Kruskal-Wallis test across all compounds to identify strains with the most divergent fitness responses between clusters (Table S10B). These “cluster-separating” strains serve as global discriminators, distinguishing compound groups based on population-wide response variation. Interestingly, 39 of the 50 strongest separating strains (78%) belonged to subpopulations enriched in the subpopulation enrichment analysis, confirming that the subpopulation-level enrichments reflect consistent, strain-specific biases rather than random effects.

Together, these results demonstrate that the fitness responses from a barcoded strain collection enable a powerful profiling framework, organizing a diverse natural compound library into functional clusters that reflect shared modes of action and reproducible, population-structured axes of sensitivity.

### Genetic associations reflect phenotypic variance across compound responses

As shown in our proof-of-concept experiments, the broad genetic diversity in our barcoded natural population allows for genome-wide association studies that links genomic variants to phenotypes. This could be particularly powerful in the context of natural compound libraries where the biological target of the compounds was largely unknown. To bridge the fitness response in the barcoded population and the potential biological target of the chemical compounds, we performed genome-wide association analysis across all tested conditions. We identified 83 significant SNP-trait associations (p < 1×10^−5^) across 107 compounds (Figure S7; Table S11A-B). These SNPs mapped to 46 genes involved in diverse processes, including genome maintenance, ribosome biogenesis, vesicular trafficking, amino acid biosynthesis and stress response processes.

These compound associations were distributed across all six fitness-based functional clusters, indicating that genetic associations span multiple response types rather than being confined to a single phenotypic class. To assess whether genetic associations were concentrated within specific compound clusters, we compared the fraction of these associated compounds across all six clusters. GWAS hit enrichment was strongest in clusters 1 and 4 (Fisher’s exact test, p = 0.03–0.04). Cluster 2 showed a mild depletion in GWAS hits (p = 0.04), whereas the remaining clusters showed no significant deviation (p > 0.05). Among the clusters, cluster 1 and cluster 4 were characterized by moderate fitness inhibition and increased phenotypic variance, whereas cluster 2 showed low fitness inhibition and relatively uniform responses (Tables S6, S9). Cluster 6 comprised only of four compounds but was defined by strong fitness inhibition and very high phenotypic variance but probably lacked detection power (p = 0.28).

Together, these analyses suggest that genetic associations are not restricted to a single compound-response class but tend to emerge under conditions of measurable fitness inhibition and increased phenotypic variability. Based on the associations, we could further classify the compound clusters into broad functional themes: Cluster 1 was linked to genome maintenance and cell-cycle progression, Cluster 2 was dominated by vesicular transport and ribosome biogenesis, Cluster 3 and 6 encompassed membrane and mitochondrial processes, Cluster 4 primarily involved genome maintenance and stress response pathways, and Cluster 5 combined metabolic regulation (Table S11A-B). These categories highlight the multifaceted physiological bases of compound responses, although the limited number of genes per cluster prevented formal enrichment.

## Discussion

Understanding how natural genetic variation shapes fitness across environments remains a central challenge in evolutionary and functional genomics ^4,58,59^. We developed a systematically barcoded collection of 520 natural *Saccharomyces cerevisiae* isolates, which captures the genetic and ecological diversity of the original 1,011-isolate population ^17^. This resource enables high-throughput, low-volume phenotyping and expands the accessible genotype-phenotype landscape of natural diversity. By integrating barcode sequencing, fitness-based clustering, and GWAS, it captures both global patterns of strain fitness and the genetic variants underlying phenotypes. Beyond environments with known functions, the barcoded population can profile compounds with unknown biological targets, as shown by GWAS linking 107 compounds to candidate genetic variants.

Systematic genetic resources have been instrumental in linking genotype to phenotype and in uncovering cellular responses to chemical perturbations. The yeast deletion collection has long served as a hallmark resource for functional profiling of chemical compounds ^37,40,54–56^. In comparison, the barcoded natural population exhibits a broader and more diverse range of bioactivities, yielding a more polarized and variable fitness landscape across nearly all shared compounds. While the deletion mutants link gene function to compound response, the barcoded natural collection reveals how naturally occurring genetic variation shapes these interactions across diverse backgrounds.

Profiling 633 natural compounds from a chemical library identified 455 that produced measurable, strain-level fitness variation across the population. These compounds grouped into six functional clusters with coherent fitness signatures, each displaying distinct subpopulation-specific patterns ^17^. For example, Ale beer and French dairy isolates consistently showed low fitness, whereas Mexican agave, West African cocoa, and Ecuadorian strains exhibited high fitness across several clusters. These structured sensitivity profiles indicate that population structure influences compound tolerance. Moreover, the observed patterns provide an evolutionary context for interpreting compound responses, suggesting that fitness variation reflects underlying adaptation to distinct ecological niches ^30,44,60,61^.

Coupling these phenotypic profiles with GWAS identified 83 significant SNP associations across 107 compounds and 46 genes involved in genome maintenance, vesicle trafficking, and ribosome biogenesis. These associations reveal genetic factors that contribute to variation in compound response, including potential links between naturally occurring polymorphisms and specific cellular processes. Although GWAS can capture a wide range of natural variants beyond loss of function, its power remains limited by allele frequency, sample size, and population structure ^62–64^. Rare variants or associations confined to small subpopulations, such as the Ale beer, French dairy, or Mexican agave groups enriched in specific compound clusters, may be underrepresented ^52,65,66^. In addition, small-effect variants underlying highly polygenic traits often fall below statistical detection thresholds, further limiting resolution for complex phenotypes ^35,65,67^.

The barcoded collection extends chemical-genetic profiling to a genetically diverse population. Similar to deletion mutant pools used for functional annotation in the reference BY background, the subpopulation-specific patterns identified here suggest that a small set of strategically chosen natural isolates could serve as representative reference panels for rapid and informative profiling of compound activity ^40,54,55^. Combining the strengths of both systems offers a powerful route to mechanistic understanding. Highly responsive natural backgrounds identified through population-level screening could serve as anchors for constructing targeted deletion libraries ^68,69^. These background-specific resources would improve precision and yield more relevant insights into compound targets and tolerance mechanisms. Expanding this approach to include time-resolved measurements, metabolic profiling, and integration with other multi-omic layers will further advance causal and mechanistic understanding. Extending beyond chemical-genetic applications, this collection has also been used to track evolutionary trajectories in mixed populations, demonstrating that initial fitness predicts long-term persistence and that adaptive changes depend on both genotype and environment ^70^.

Overall, the barcoded collection offers a scalable framework to connect natural genetic diversity with phenotypic variation, bridging evolutionary and functional genomics at population scale.

## Materials and methods

### Strains used

The *S. cerevisiae* strains used in this study are described in Table S1 as per 1,011 *S. cerevisiae* collection ^17^. Strains were either grown on YPD (yeast extract 1%, peptone 2%, dextrose 2%) medium or YPD medium supplemented with G418 (Euromedex – 4ml/l of 50 mg/ml stock) and hygromycin (Jena Bioscience – 6 ml/l of 50 mg/ml stock) at 30°C ^71,72^. All strains are available upon request.

### Barcoding of the 1,011 isolates

We generated a systematically barcoded collection of *Saccharomyces cerevisiae* natural isolates using a PCR-mediated gene disruption strategy ^17^. A barcode integration cassette was designed for targeted insertion at the neutral *HO* locus via homologous recombination. The construct contained a selectable marker, either *KanMX* or *HphMX*, conferring resistance to geneticin or hygromycin, respectively, flanked by unique 20-nucleotide barcode sequences serving as molecular identifiers for individual strains. Barcode cassettes were generated by PCR amplification of the *KanMX/HphMX* resistance modules using synthetic oligonucleotides (Eurofins Genomics GmbH) containing 50 bp homology arms to the *HO* locus and 20 randomized nucleotides to encode unique barcode identifiers. The forward primer (BCF) and reverse primer (BCR) amplified the full barcode marker cassette (Figure 1A). All primer sequences used in the study are provided in Table S12. The pooled barcode constructs (~0.5 μg/μL) were independently transformed into each isolate from the 1,011-strain *S. cerevisiae* collection using a lithium acetate transformation protocol adapted to a 96-well plate format ^73^. Up to five transformation rounds were performed to maximize integration efficiency. Transformants were selected on antibiotic-containing media and verified for correct barcode insertion by diagnostic PCR targeting the *HO* locus. Barcode sequences were determined by Sanger sequencing of the forward barcode region. Eventually, we were able to barcode, sequence and verify 520/1,011 isolates.

### Pooled Fitness Measurement Assay (Bar-Seq)

Equal volumes of independent saturated cultures of the 495 barcoded isolates were pooled after 48 hours. This pooled cultured formed the initial 0 hour timepoint. Initial pool was inoculated into 200 uL YPD media supplemented with compounds or YP-galactose/ glycerol (Tables S2, S4) at an equivalent starting OD_600_ of 0.05 (~10^6^ cells per ml). The pooled cultures were propagated at 30 degrees and harvested at 24 hour timepoint for DNA extraction and sequencing.

### DNA extractions, PCR amplification of the barcode locus, multiplexing and sequencing

Briefly, harvested cultures were resuspended in 200mM lithium acetate + 1% SDS solution and incubated at 70 degrees for 15 mins. DNA was precipitated by treating with 100% ethanol at −20 degrees. The samples were then subjected to high-speed centrifugation and washed with 70% ethanol. The pellet was resuspended in 50 uL of 0.1X TE buffer. This was subsequently subjected to high-speed centrifugation, and the supernatant was collected for PCR amplification of the barcode locus. We employed a two-step PCR approach to amplify the barcodes from the DNA utilizing the Q5-hi fidelity polymerase kit from NEB, following a protocol as described in Levy et al. 2015 ^33^. The primer sequences and conditions used are listed in Table S12. We used the Qubit HS kits to assess the concentration of the PCR product for each of the samples. Subsequently, we combined them in equimolar ratios into a single sample for high-throughput Illumina sequencing at Novogene Europe.

### Initial processing of the amplicon sequence data

The paired reads were first merged with fastp, before being converted from fastq to fasta format with seqtk (). A blast database was generated from all these files and queried with the blastn program from a fasta file containing 538 non-ambiguous barcodes related to strains with the following parameters: - word_size 13; -strand plus; -dust no (Table S1) ^74^. Number of occurrence of each barcode was extracted from the blast result files.

### Genome-wide association analysis

Genome-wide association studies (GWAS) were performed to identify genetic variants associated with variation in compound-specific fitness phenotypes across the barcoded *S. cerevisiae* collection. We used a high-density genotype matrix comprising 91,317 single nucleotide polymorphisms (SNPs) and 700 copy number variants (CNVs), corresponding to an average marker density of approximately one variant per 100 bp. SNP genotypes were derived from the *S. cerevisiae* 1,011 genomes dataset (and filtered using bcftools v1.16 (with the following parameters: minimum read depth (DP) ≥ 10, genotype quality (GQ) ≥ 20, ≥ 99% non-missing data per site, and restriction to biallelic variants ^17,75^. CNVs were quantified for each gene in the pangenome using normalized read depth, calculated as the ratio between the median coverage depth of a gene and the median coverage depth of the whole genome. For each gene, CNVs were binarized across multiple normalized depth thresholds (0.25 to 2.25, in increments of 0.5), reflecting expected copy-number variation in predominantly diploid isolates. A combined PLINK-compatible genotype matrix was generated by treating each gene-threshold combination as an independent variant, encoding genotypes as “1” or “2” for values above or below the respective threshold. SNP and CNV matrices were merged, filtered for minor allele frequency (MAF) ≥ 5%, and linkage disequilibrium (LD) pruned using PLINK v1.9 with parameters --maf 0.05 and --indep-pairwise 50kb 1 0.5 ^76^. The resulting combined matrix served as the final genotype dataset for GWAS. Phenotypic traits were defined as compound-specific fitness scores (log_2_ fold change values) from pooled competition assays. Each phenotype was normalized using a rank-based inverse normal transformation to reduce the influence of outliers and non-normal trait distributions. Associations were tested using a linear mixed model (LMM) implemented in FaST-LMM v0.6.4, including ploidy as a fixed covariate to account for its known effects on phenotypic variance ^77^. A permutation-based significance threshold (100 permutations per trait; α = 0.05) was used to control for multiple testing and define genome-wide significance.

### Compound similarity matrix and hierarchical clustering

To identify global structure in compound responses, we computed pairwise correlations between all compounds based on their strain-level fitness profiles. Fitness values were derived from the normalized log_2_ fold change (log_2_FC) of each strain relative to baseline abundance across all compounds. For each pair of compounds, a Pearson correlation coefficient (r) was calculated across all strains, producing a symmetric compound–compound similarity matrix. This matrix captured the degree of similarity in strain-level response profiles, where higher correlations indicate similar phenotypic signatures and potentially shared modes of action.

The resulting similarity matrix was converted into a distance matrix (1 − r) and used as input for hierarchical clustering. Clustering was performed using the Ward’s minimum variance method (ward.D2 linkage) implemented in R’s hclust() function. The optimal number of clusters (k = 6) was determined using the elbow method, where the total within-cluster variance was plotted against the number of clusters, and the inflection point corresponding to the largest gain in explanatory power was selected. Cluster assignments were extracted from the dendrogram cut and appended to the metadata. Each cluster thus represents a distinct group of compounds with correlated strain-level fitness profiles, reflecting shared bioactivity patterns or related mechanisms of action.

### Subpopulation-based enrichment analysis

To identify population-level signatures of compound response, we performed a gene set enrichment– like analysis to test whether specific *S. cerevisiae* subpopulations were disproportionately represented among strains showing strong fitness responses within each compound or compound cluster. Each strain in the barcoded collection was assigned to one of the major population subgroups defined previously by population structure ^17^. For each compound, we computed the mean log_2_ fold change (fitness score) for every strain relative to the population median. Strains were ranked by these fitness scores, producing a compound-specific ranked list. Subpopulation enrichment was quantified using a GSEA-like running-sum statistic, comparing the observed distribution of subpopulation members across the ranked strain list against a random expectation derived from 10,000 phenotype label permutations ^57^. The enrichment score (ES) reflects the degree to which strains from a given subpopulation are overrepresented among the most sensitive or resistant individuals. Normalized enrichment scores (NES) and associated empirical P-values were calculated using the permutation-based null distribution.

To summarize global patterns, enrichment scores were averaged within each of the six compound clusters defined by compound–compound similarity analysis. False discovery rate (FDR) correction was applied across all subpopulation–compound pairs using the Benjamini–Hochberg method. Significant enrichment was defined as |NES| > 1 and FDR < 0.05.

## Data availability

The barcoded strain collection is available upon request. Sequence data are available from National Centre for Biotechnology Information Sequence Read Archive under accession number: PRJEB96750.

## Acknowledgements

We thank the members of the Petrov lab at Stanford University for their advice on experimental design and methodology. This work was supported by the European Research Council (ERC Consolidator Grant 772505). JS is a Fellow of the University of Strasbourg Institute for Advanced Study (USIAS) and a member of the Institut Universitaire de France. We are grateful to Ms. Manami Morihashi for preparing compound samples and the Université de Strasbourg-RIKEN Researcher Exchange Program for facilitating the collaboration. This work was supported by a Grant-in-Aid for Scientific Research from the Japan Society for the Promotion of Science (25K01916 to CB and YY) and a Grant-in-Aid for Transformative Research Areas (A) “Latent Chemical Space” from the Ministry of Education, Culture, Sports, Science and Technology, Japan (23H04882 to YY).

