## Supplementary material for "Population-scale chemical response revealed by a barcoded yeast collection": Figure S1

A

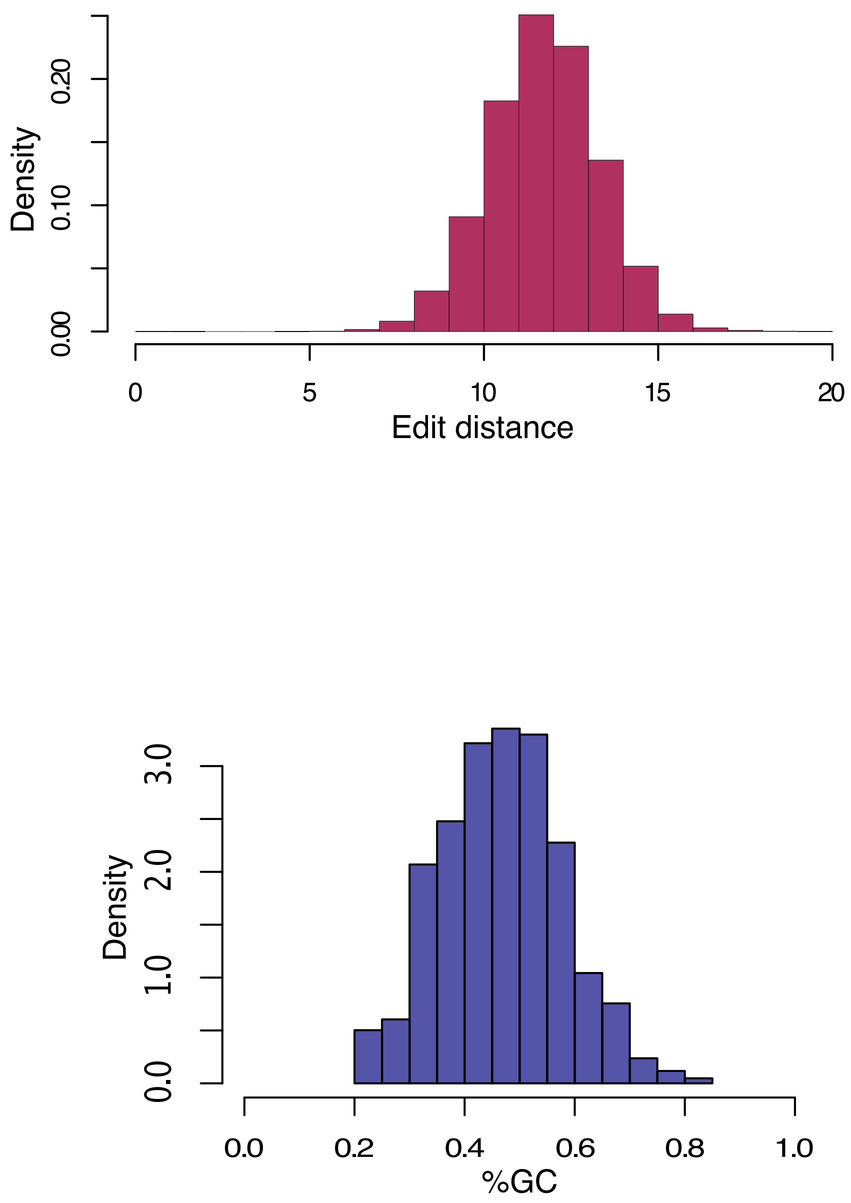


**Figure S1.** Barcode sequence composition and diversity. **A.** Distribution of Levenshtein edit distances among all barcode sequences, indicating high sequence diversity and minimal overlap between barcodes. **B.** Distribution of GC content (%) across all barcode sequences, showing balanced nucleotide composition without GC bias.

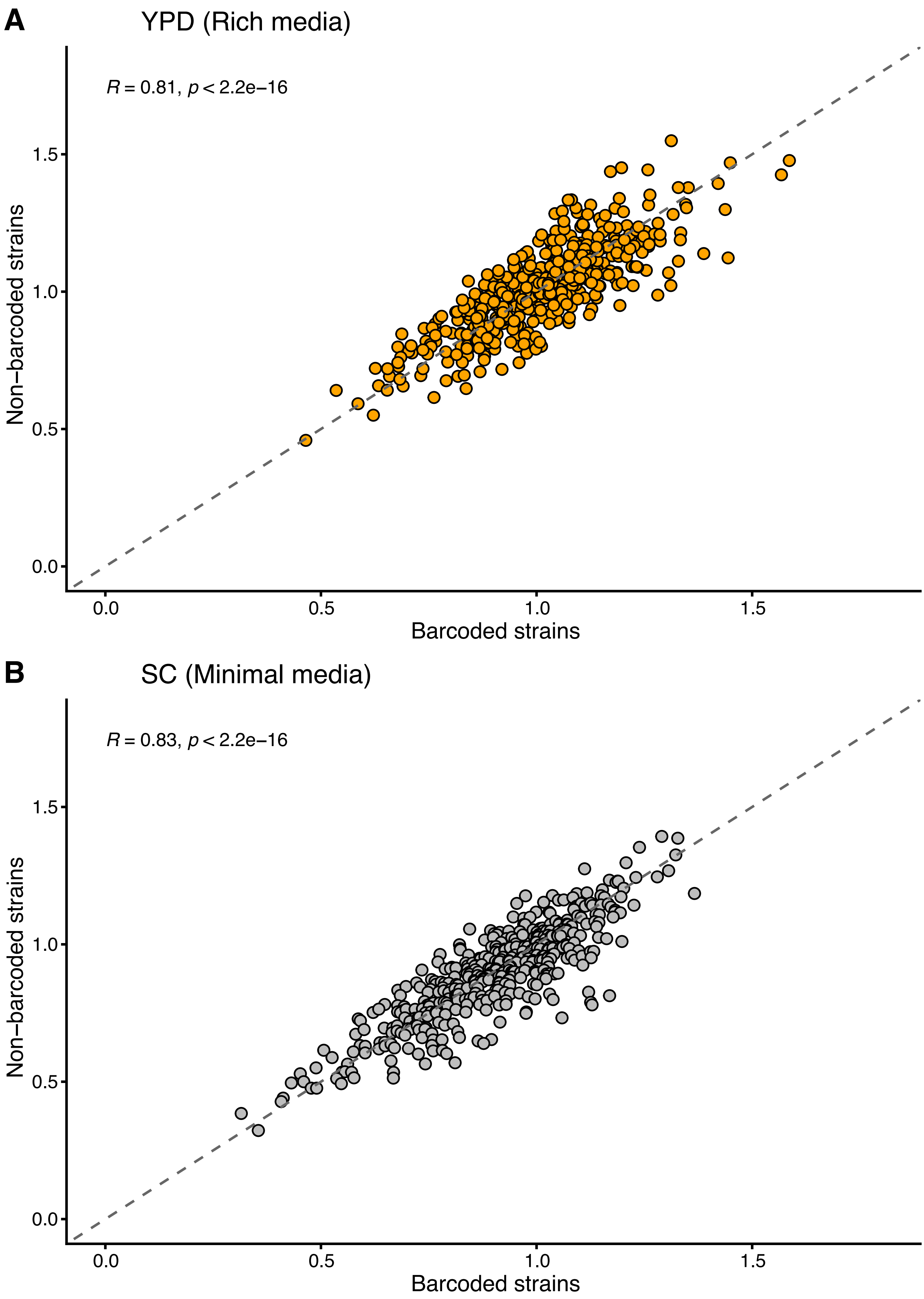


**Figure S2.** Barcoding does not alter strain fitness under standard growth conditions. **A.** Scatterplot of fitness (colony size relative to the BY reference) between barcoded and non-barcoded isolates grown in rich medium (YPD). **B.** Corresponding comparison in minimal medium (SC). High correlations ($R = 0.81$–$0.83$, $p < 2.2 \times 10^{-16}$) indicate that barcode integration does not measurably affect strain fitness in either condition.

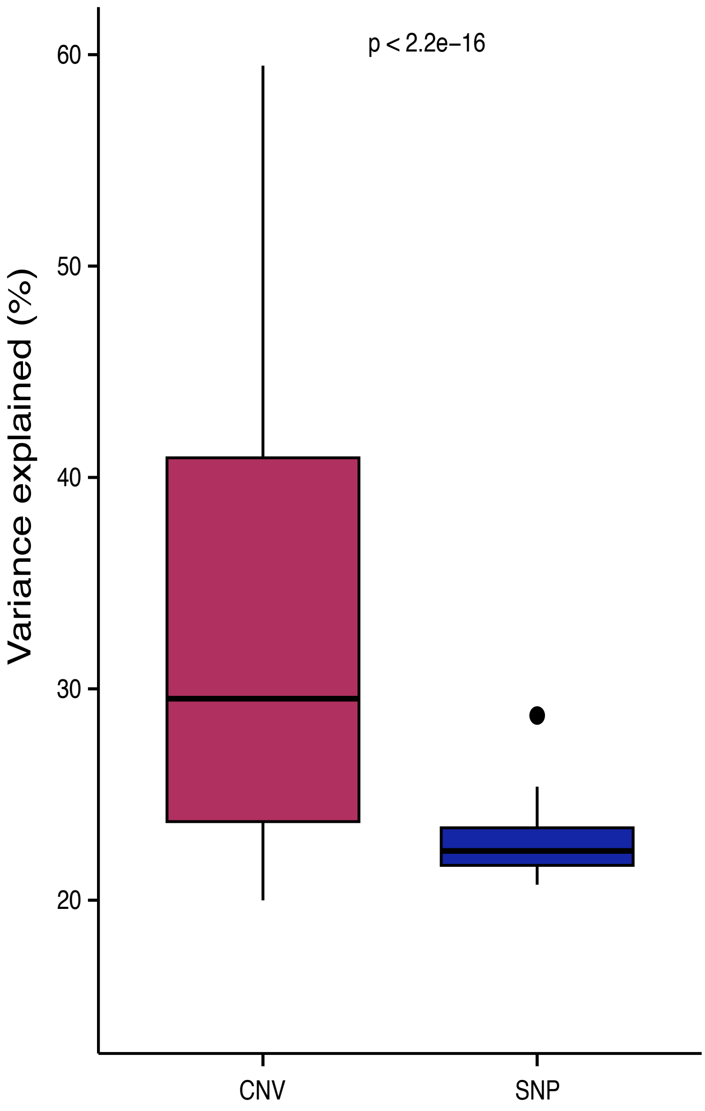


**Figure S3.** Copy number variants explain a greater proportion of fitness variance than SNPs. Boxplot showing the distribution of variance explained by significant CNV- and SNP-based associations across 13 compounds in our benchmarking experiment.

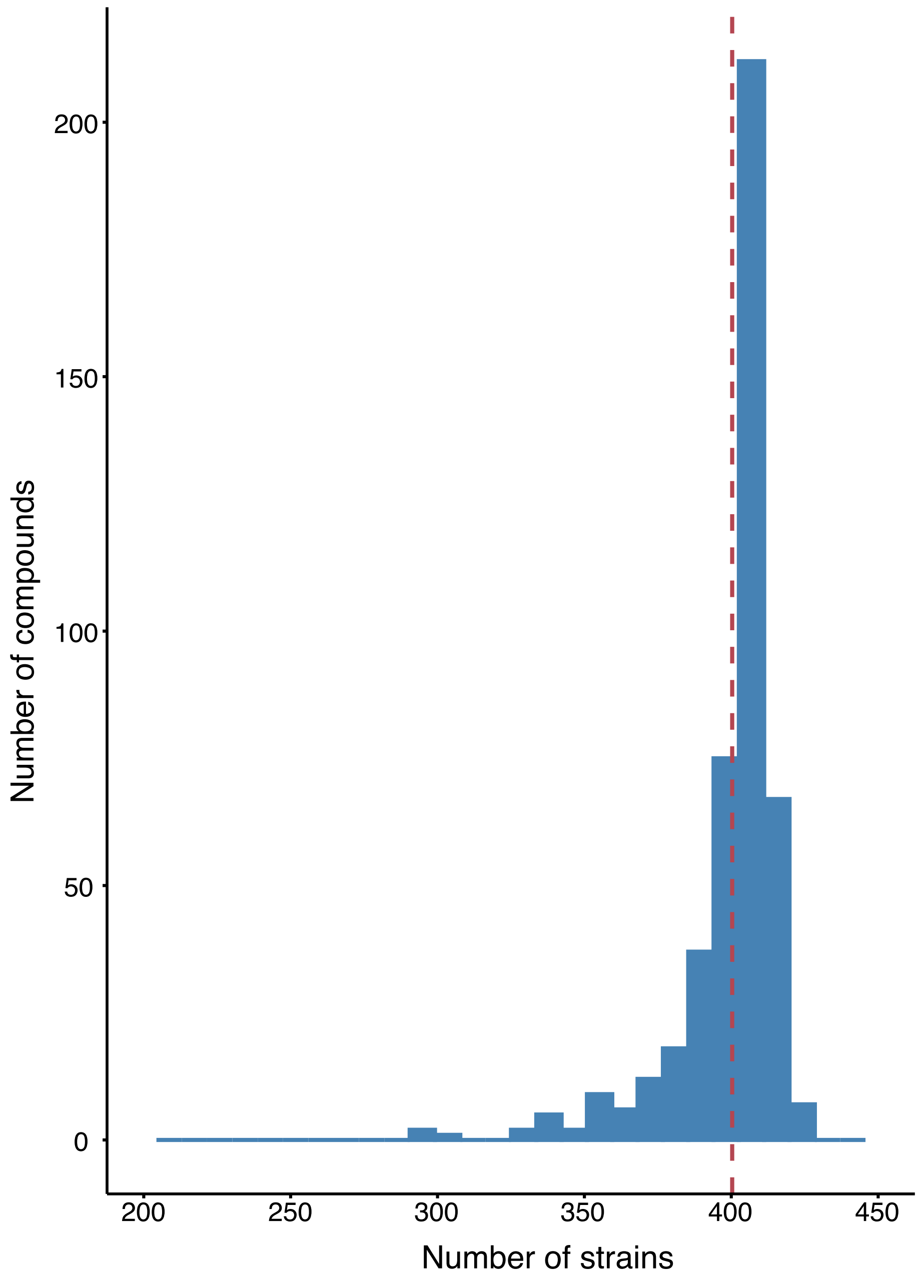


**Figure S4.** Distribution of strain detection across the compound library. Histogram showing the number of strains detected at the 24-h timepoint across 455 compounds that passed our QC threshold. On average, ~400 strains were detected per condition.

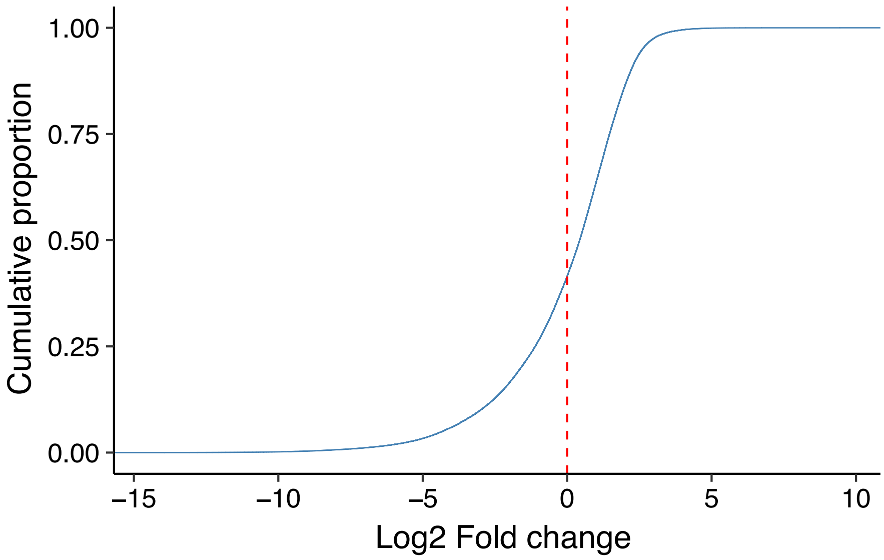


**Figure S5.** Global distribution of strain fitness across all compound treatments. Empirical cumulative distribution function (ECDF) of strain fitness values ($\log_2$ fold change) across 455 compounds. The fitness distribution spans both positive and negative values but is asymmetric relative to neutrality (0), indicating unequal representation of fitness gains and losses and suggesting that subsets of strains respond non-randomly to chemical perturbations.

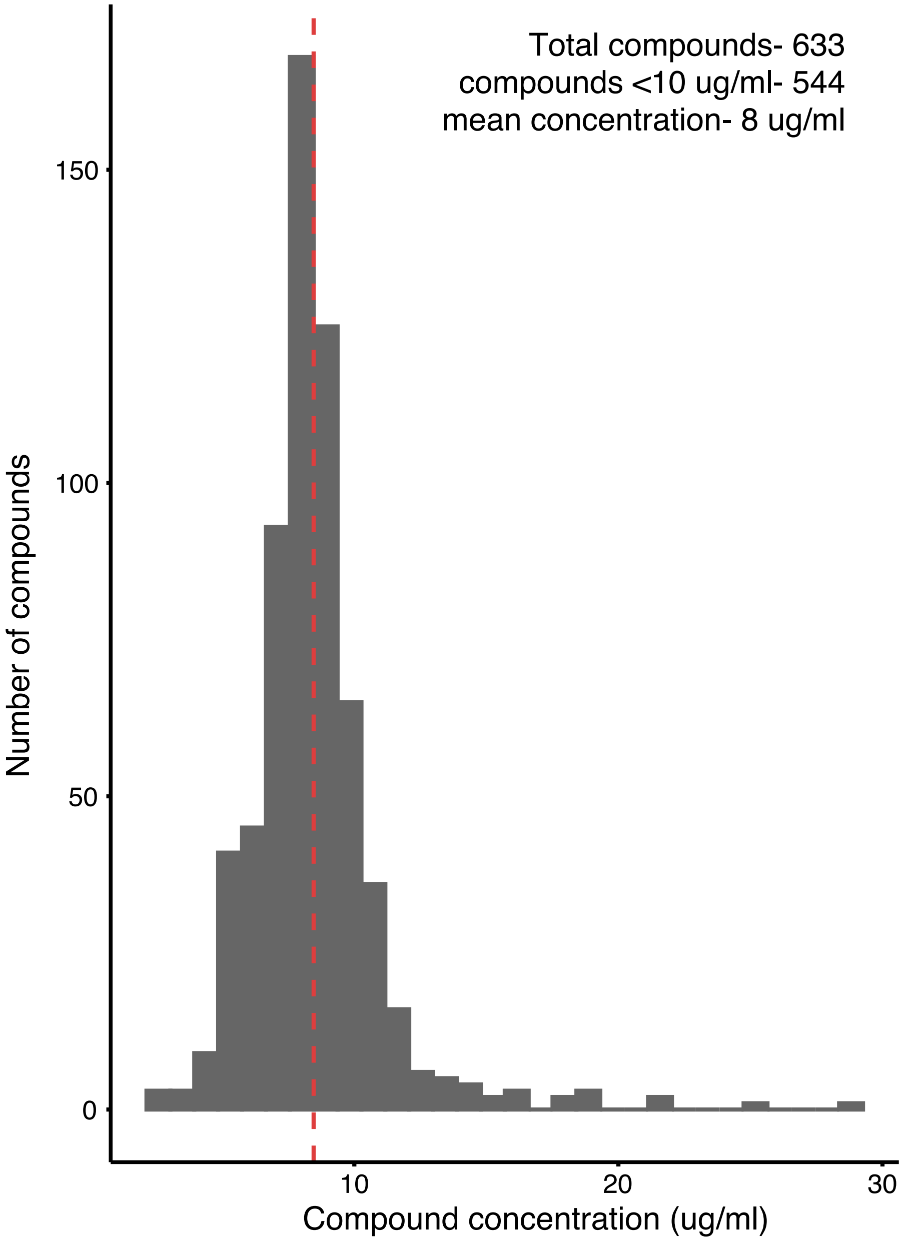


**Figure S6.** Compound concentration distribution in the barcoded collection screen. Histogram showing the concentration distribution of the 633 compounds screened in this study (mean = 8 µg/mL). Of these, 544 compounds (≈ 80%) were assayed at concentrations below 10 µg/mL. Piotrowski et al. 2017, screened at a uniform concentration of 10 µg/mL. Thus, most compounds in the present study were assayed at equal or lower effective concentrations.

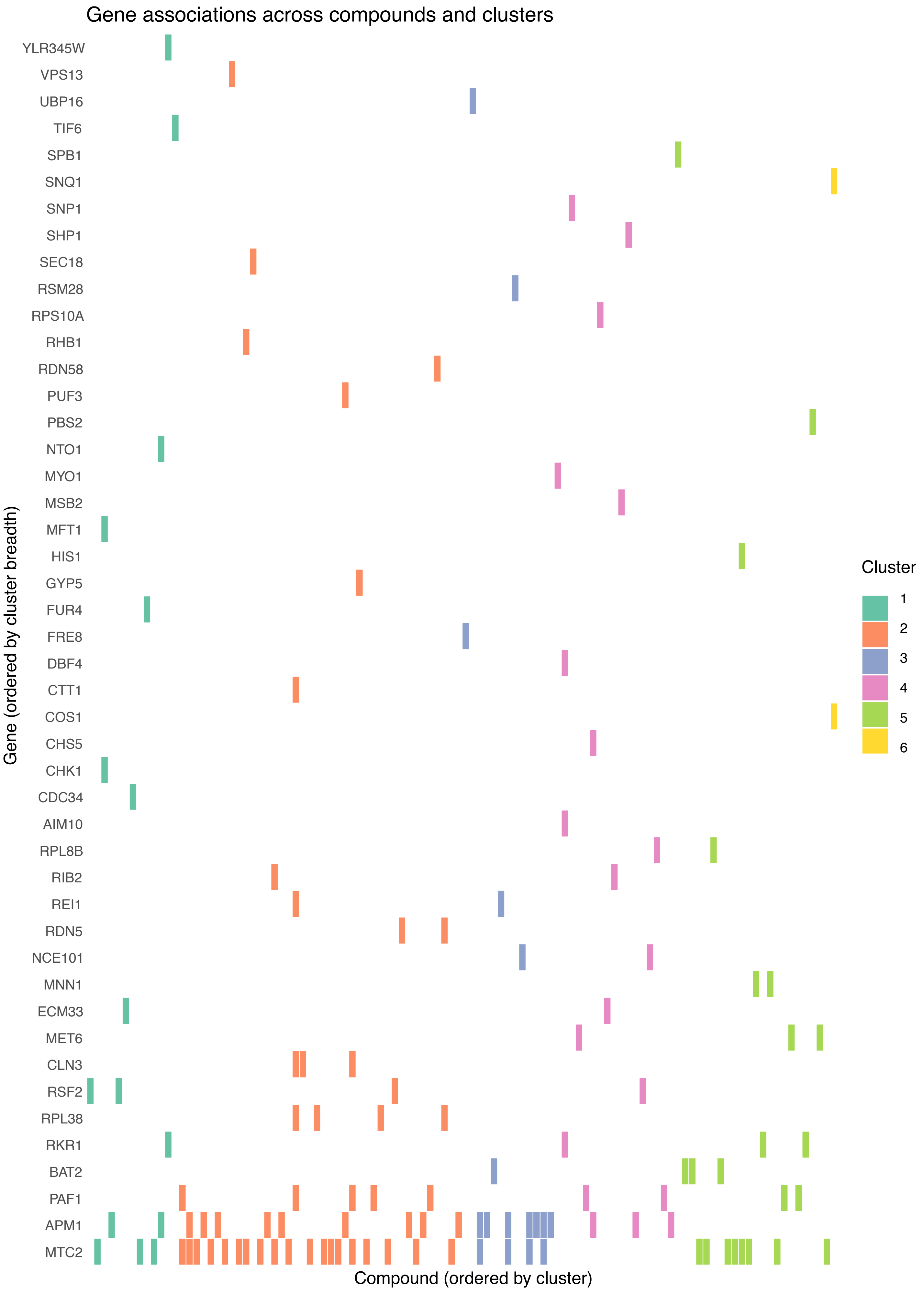


**Figure S7.** Genetic associations between SNPs and compounds. SNP-trait associations ($p < 1 \times 10^{5}$) across 107 compound conditions. In total, 83 associations mapped to 46 genes, spanning the six compound clusters.
